# Artificial cerebellum on FPGA: Realistic real-time cerebellar spiking neural network model capable of real-world adaptive motor control

**DOI:** 10.1101/2023.05.23.541669

**Authors:** Yusuke Shinji, Hirotsugu Okuno, Yutaka Hirata

**Affiliations:** Department of Computer Science, Chubu University Graduate School of Engineering, Kasugai, Japan; Faculty of Information Science and Technology, Osaka Institute of Technology, Hirakata, Japan; Department of Artificial Intelligence and Robotics and Technology, Chubu University Graduate School of Engineering, Kasugai, Japan; Center for Mathematical Science and Artificial Intelligence, Chubu University, Kasugai, Japan; Chubu University Academy of Emerging Sciences, Kasugai, Japan

**Keywords:** Artificial cerebellum, Spiking Neural Network, FPGA, Adaptive control, Motor learning

## Abstract

The cerebellum plays a central role in motor control and learning. Its neuronal network architecture, firing characteristics of component neurons, and learning rules at their synapses have been well understood in terms of anatomy and physiology. A realistic artificial cerebellum with mimetic network architecture and synaptic plasticity mechanisms may allow us to analyze cerebellar information processing in real world by applying it to adaptive control of actual machines. Several artificial cerebellums have previously been constructed, but they required a high-performance hardware to run in real time for real-world machine control. Presently, we implemented an artificial cerebellum with the size of 10^4^ spiking neuron models on a field-programmable gate array (FPGA) which is compact, lightweight, portable, and low-power-consumption. In the implementation three novel techniques are employed: 1) 16-bit fixed-point operation with randomized rounding, 2) fully connected spike information transmission, 3) alternative memory that uses pseudo-random number generators. We demonstrate that the FPGA artificial cerebellum runs in real time, and its component neuron models behave as those in the corresponding artificial cerebellum configured on a personal computer in Python. We applied the FPGA artificial cerebellum to adaptive control of a machine in real-world and demonstrate that the artificial cerebellum is capable of adaptively reducing control error after sudden load changes. This is the first implementation and demonstration of a spiking artificial cerebellum on an FPGA applicable to real-world adaptive control. The FPGA artificial cerebellum may provide neuroscientific insights into cerebellar information processing in adaptive motor control and may be applied to various neuro-devices to augment and extend human motor control capabilities.

## 1 Introduction

The cerebellum has been shown to play crucial roles in adaptive motor control of eye movements (McLaughlin, 1967; Ito et al., 1970; Miles et al., 1986; Nagao, 1988), blinks (Lincoln et al., 1982; Yeo et al., 1985), arm reaching (Martin et al., 1996; Kitazawa et al., 1998), gait (Mori et al., 1999; Ichise et al., 2000), and posture (Nashner, 1976) among others (Monzée et al., 2004; Leiner, 2010). It has been envisioned that if the cerebellar function is realized artificially as a realistic artificial cerebellum, it could be used to understand neural mechanisms of cerebellar motor learning as well as to control actual machines such as robots and drones in real-world.

Cerebellar neural circuitry has been well understood in terms of anatomical connectivity and physiological neuronal characteristics (Eccles et al., 1967; Gao et al., 2012). Several artificial cerebellums have been constructed since the pioneering theoretical work by Marr (Marr, 1969) and Albus (Albus, 1971). These models can be categorized into two major types, spiking and non- spiking, depending on the type of neuron model conforming the cerebellar network. In general, non- spiking models require much less computational power than spiking models, making it possible to simulate a larger scale model in real-time (Pinzon-Morales and Hirata, 2014, 2015; Vijayan and Diwakar, 2022). However, non-spiking models have a limitation in addressing computational roles of spike timings in cerebellar information processing, especially those of synchronous spiking of Purkinje cells (Welsh et al., 1995; Sedaghat-Nejad et al., 2022) as well as spike timing dependent synaptic plasticity (STDP) that were found to be a fundamental neural basis of cerebellar motor learning (Ito et al., 1982; Hirano, 1990; De Zeeuw et al., 2021). Spiking models are necessary to study these aspects of cerebellar information processing. A major caveat of spiking models in oppose to non-spiking ones is their computational costs. For example, an artificial cerebellum with 68 billion neurons required a supercomputer (K computer, RIKEN Center for Computational Science) to be executed in real time (Yamaura et al., 2020) while it took 30 times more than real-time to run an artificial cerebellum with a size of 10 thousands neurons on an ordinary personal computer (Inagaki and Hirata, 2017), which would be 5 times even on a latest high spec personal computer (Intel i9- 7900x CPU, 128 GB memory).

Field-programmable gate arrays (FPGAs) are known to excel in parallel computing, thus may be suitable for implementation of a spiking artificial cerebellum to be executed in real time. Luo et al (Luo et al., 2016) and Xu et al. (Xu et al., 2018) have successfully implemented an artificial cerebellum with 10^4^ spiking neurons on an FPGA (Xilinx Kintex-7 KC705), and applied it to neuro- prosthesis in rats, using an eye blink conditioning scheme. However, this first FPGA spiking artificial cerebellum uses 40-bit fixed-point operation that consumes a large number of arithmetic and storage devices on the FPGA. It also stores each and every connection information between numerous neuron models, resulting in occupying a large portion of the storage devices. Furthermore, it has a communication delay which depends on the number of total spikes in the artificial cerebellum, resulting in slower execution when more spikes are generated in the model. Due to these configurations, molecular layer inhibitory interneurons (basket/stellate cells) were not implemented in the previous artificial cerebellum.

Presently, we implemented an artificial cerebellum on an FPGA composed of major cerebellar cortical neuron types including molecular layer inhibitory interneurons. We employed three novel implementation strategies to achieve real-time execution of the artificial cerebellum with the scale of 10^4^ neurons: First, we reduced the required number of arithmetic and storage devices while maintaining numerical accuracy by using only 16-bit fixed-point numbers and introducing randomized rounding when computing numerical solutions to the differential equations describing each spiking neuron model. Second, when spike information is transmitted between the computational units of each neuron, we installed a spike storage unit with a data transmission circuit that is fully coupled between the pre- and postsynaptic neurons, thereby eliminating the latency dependent on the number of neurons and spikes. Third, we reduced the required number of storage devices by introducing a pseudo-random number generator as a storage device for storing information about connections between neuron models.

We performed two kinds of evaluations of the FPGA spiking artificial cerebellum to evaluate its validity. In the first evaluation, we compare firing properties of a minimum scale cerebellar neuron network model implemented on the FPGA with the same model implemented on a personal computer in Python using 64-bit floating point number to demonstrate sufficient computational accuracy on the FPGA to simulate spiking timings. In the second evaluation, we apply the FPGA spiking artificial cerebellum to real-world adaptive control of a direct current (DC) motor and show that the artificial cerebellum is capable of adaptively controlling the DC motor even when its load is suddenly changed under the noisy natural environment.

## 2 Materials and Methods

### 2.1 Cerebellar spiking neuronal network model

The artificial cerebellum to be implemented on the FPGA in the present study is similar to those previously constructed by referring to anatomical and physiological evidence of the cerebellar cortex (Medina et al., 2000; Inagaki and Hirata, 2017; Casali et al., 2019; Kuriyama et al., 2021). Presently, the scale of the artificial cerebellum (the number of neuron models) is set to ∼10^4^ neurons which is limited by the specification of the FPGA (XC6SLX100, Xilinx) used in the current study (see below for more detailed specs). The size is easily expandable by connecting other FPGAs.

#### 2.1.1 Network structure

The artificial cerebellum has a bi-hemispheric structure as the real cerebellum (Figure 1). Each hemisphere consists of 246 mossy fibers (MF), 8 climbing fibers (CF), 4096 granule cells (GrC), 369 Golgi cells (GoC), 25 molecular layer inhibitory interneurons (MLI), and 8 Purkinje cells (PkC). These numbers were determined so that the model can be implemented and run in real time on the FPGA currently employed (see below) while preserving convergence-divergence ratios of these fiber/neuron types (Figure 1, purple numbers) as close as those found in the vertebrate cerebellum (Medina et al., 2000). Note that this number of GrC (4096) has been demonstrated to be enough to control real-world objects such as a two-wheeled balancing robot robustly (i.e., independently on the initial synaptic weight values) (Pinzon-Morales and Hirata, 2015). In the cerebellar cortical neuronal network, MFs connect to GrCs and GoCs via excitatory synapses, and GrCs connect to PkCs, GoCs, and MLIs via excitatory synapses. GoCs connect back to GrCs via inhibitory synapses, while MLIs connect to PkCs via inhibitory synapses. PkCs connect to the extracerebellar area via inhibitory synapses.

**Figure 1.**
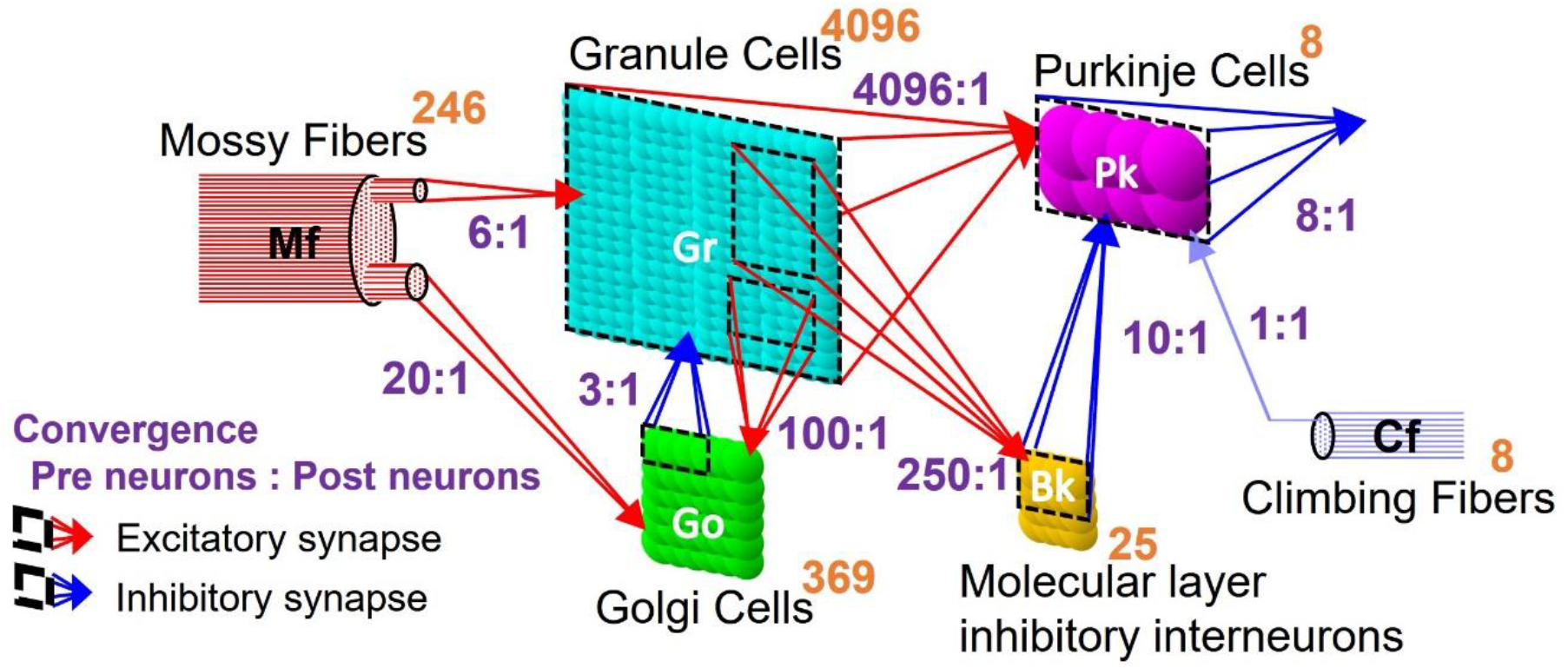
The structure of an artificial cerebellum. The number in the upper right of the neuron indicates the number of neurons. The neuron connection ratio indicates the ratio of presynaptic neurons to postsynaptic neurons, which is a convergence. Ratio of numbers of different neuron types are indicated in orange numbers. Input-output ratio of synaptic connections are indicated in purple numbers.

#### 2.1.2 Spiking neuron model

Neurons and input fibers in the artificial cerebellum are described by the following leaky integrate and fire model (Gerstner and Kistler, 2002; Izhikevich, 2010):

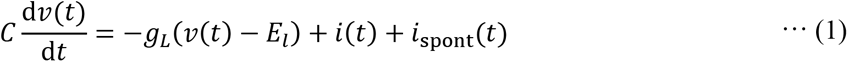

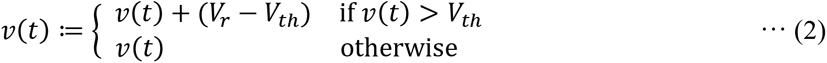

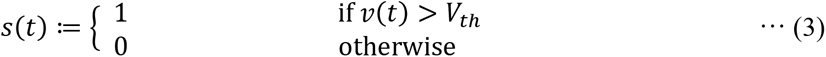

where, *ν*(*t*), *i*(*t*), and *i*_spont_(*t*) are the membrane potential, the input synaptic current, and the current producing spontaneous firing at time *t*, respectively. *C* is the membrane capacitance, and *g_L_* is the leak conductance. When the membrane potential *ν*(*t*) exceeds the threshold *Vth*, it is reset to the resting potential *V_r_* and the spike event function *s*(*t*) outputs 1. *i*_spont_(*t*) is a uniform random number [0, *I_spont_*] to describe a dc firing rate of each neuron. The constants for each neuron/fiber type are listed in Table 1 which are the same as the previous realistic artificial cerebellums (Casali et al., 2019; Kuriyama et al., 2021) except that the parameters of the input fibers were arbitrarily defined so that their firing frequencies become physiologically appropriate.

#### 2.1.3 Synapse model

The synaptic transmission properties are described by the following conductance-based synapse model (Gerstner and Kistler, 2002; Izhikevich, 2010):

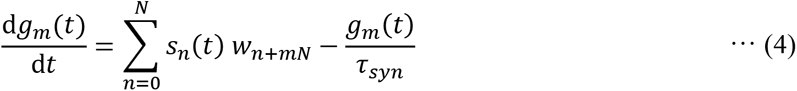

**Table 1.** Neuron parameters

| Cell type | $C$<br>[pF] | $g_L$<br>[mS] | $E_L$<br>[mV] | $I_{spont}$<br>[pA] | $V_r$<br>[mV] | $V_{th}$<br>[mV] |
| --- | --- | --- | --- | --- | --- | --- |
| MF | 1.0 | 0.03 | -70 | 5.0 | -80 | -55 |
| CF | 1.0 | 0.3 | -70 | 0.015 | -80 | -55 |
| GrC | 3.0 | 1.5 | -65 | 0.0 | -75 | -55 |
| GoC | 76.0 | 3.6 | -74 | 36.8 | -84 | -42 |
| BkC | 14.6 | 1.0 | -68 | 15.6 | -78 | -53 |
| PkC | 620.0 | 7.0 | -62 | 600.0 | -72 | -47 |

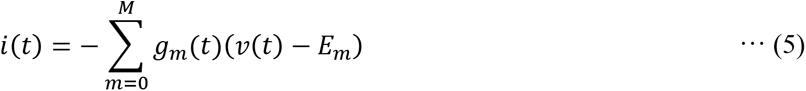

where, *g_m_*(*t*) is the synaptic conductance of the *m*-th postsynaptic neuron, *s_n_*(*t*) is the spike event function of the *n*-th presynaptic neuron or fiber, and *w_n+mN_* is the synaptic transmission efficiency between the *n*-th presynaptic neuron or fiber and the postsynaptic neuron. *N* is the number of presynaptic neurons. *τ_sym_* is the time constant. *E_m_* is the reversal membrane potential which is positive or negative for excitatory or inhibitory synapse, respectively. As a result, the sign of the synaptic current *i*(*t*) differs between the excitatory synapse and the inhibitory synapse. *M* is the number of presynaptic neuron types. The ratios of the numbers of synaptic connections between different neuron types were as shown in Figure 1 (N_c1_:N_c2_) where N_c1_ and N_c2_ are the numbers of presynaptic and postsynaptic neurons, respectively. The connections between presynaptic and postsynaptic neurons are determined by a pseudo-random number generator described later. The initial values of synaptic transmission efficiency *w* of all synapses were assigned by Gaussian random numbers whose means and variances are different for different neuron types as listed in Table 2. In the current model, only parallel fiber (PF, the axonal extensions of GrC) - PkC synapses undergo synaptic plasticity as described below. Other synaptic efficacies were fixed at the initial value throughout the execution. The synaptic constants *τ_sym_* and *E_m_* were set as shown in Table 2 based on anatomical and physiological findings (Medina et al., 2000; Kuriyama et al., 2021).

**Table 2.** Synapse parameters

| Postsynaptic<br>neuron | Presynaptic<br>neuron | Mean of<br>initial weight $w$ [pS] | Reversal potential<br>$E$ [mV] | Time constant<br>$\tau_{syn}$ [ms] |
| --- | --- | --- | --- | --- |
| GrC | MF | 19.5 | 0 | 0.50 |
| GrC | GoC | 3.9 | -70 | 10.00 |
| GoC | MF | 171.9 | 0 | 0.50 |
| GoC | GrC | 31.2 | 0 | 0.50 |
| PkC | GrC | 0.0 | 0 | 0.50 |
| PkC | BkC | 3.9 | -70 | 1.60 |
| BkC | GrC | 3.9 | 0 | 0.64 |

#### 2.1.4 Synaptic plasticity

The synapses between the PF and the PkC are the loci where the memory of motor learning has been proposed to be stored (Ito, 2001; Takeuchi et al., 2008). These include long-term depression (LTD) and long-term potentiation (LTP). The present model implements plasticity as follows:

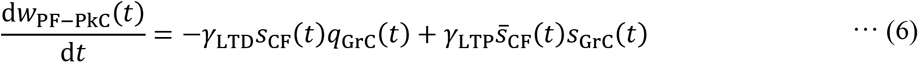

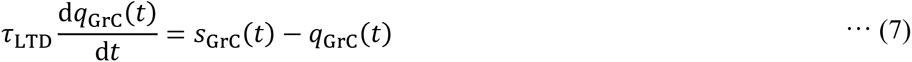

Here, *q*_GrC_ represents the average firing rate of the GrC, *s*_GrC_(*t*) represents a spike event of the GrC, and *s̄*_CF_(*t*) represents the negation of a spike event of the CF. *s*_GrC_(*t*) turns to 1 when a spike fires, otherwise 0. The synaptic weight *w*_PF–PkC_(*t*) increases or decreases from the initial value 0 in range of [0, 1]. When the firing of GrC and CF are synchronized, LTD occurs (Ekerot and Jömtell, 2003). This plasticity model is achieved by reducing the synaptic transmission efficiency *w*_PF–PkC_(*t*) by the product of *q*_GrC_(*t*) and the coefficient *γ*_LTD_ = 5.94×10^−8^ when CF spikes at time *t*. The average firing rate *q*_GrC_ is described by the low-pass filter which have the time constant (τ_LTD_ = 100 ms). On the other hand, LTP is induced when GrC fires and CF does not fire (Hirano, 1990; Jörntell and Hansel, 2006). This plasticity model is achieved by increasing the synaptic transmission efficiency *w*_PF–PkC_(*t*) by the product of *s*_CF_(*t*), *s*_GrC_(*t*), and the coefficient *γ_LTP_* = 4.17×10^−7^.

### 2.2 FPGA implementation

#### 2.2.1 Design of motor control system

The artificial cerebellum was implemented on an FPGA (XC6SLX100-2FGG484C, Xilinx) attached to an evaluation board (XCM-018-LX100, Humandata). We used a hardware description language, VHDL to describe the artificial cerebellum which is available at [https://github.com/ to be uploaded when accepted]. The left and right hemisphere models were combined as shown in Figure 2 to control real-world machines. A proportional-differential (PD) controller that simulates the pathway outside the cerebellum was implemented on the same FPGA as in the previous models (Pinzon- Morales and Hirata, 2014, 2015). The sum of the left and right hemisphere outputs and the output of the PD controller drive control objects. The control object currently tested is a DC motor (JGA25- 370, Open Impulse). As a load added to the control object, the same type of DC motor was connected co-axially. The load was imposed by short-circuiting the DC motor via a relay circuit controlled by the same FPGA. Produced motion of the control object in response to a given target speed was measured by an encoder and fed-back through a hole sensor to calculate the error. The error is sent to the artificial cerebellum as CF activity which induces PF–PkC synaptic plasticity (see 2.4). Other input modalities to the artificial cerebellum via MFs are target speed, error, and the copy of motor command (efference copy) as in the real oculomotor control system (Noda, 1986; Hirata and Highstein, 2001; Blazquez et al., 2003; Huang et al., 2013).

**Figure 2.**
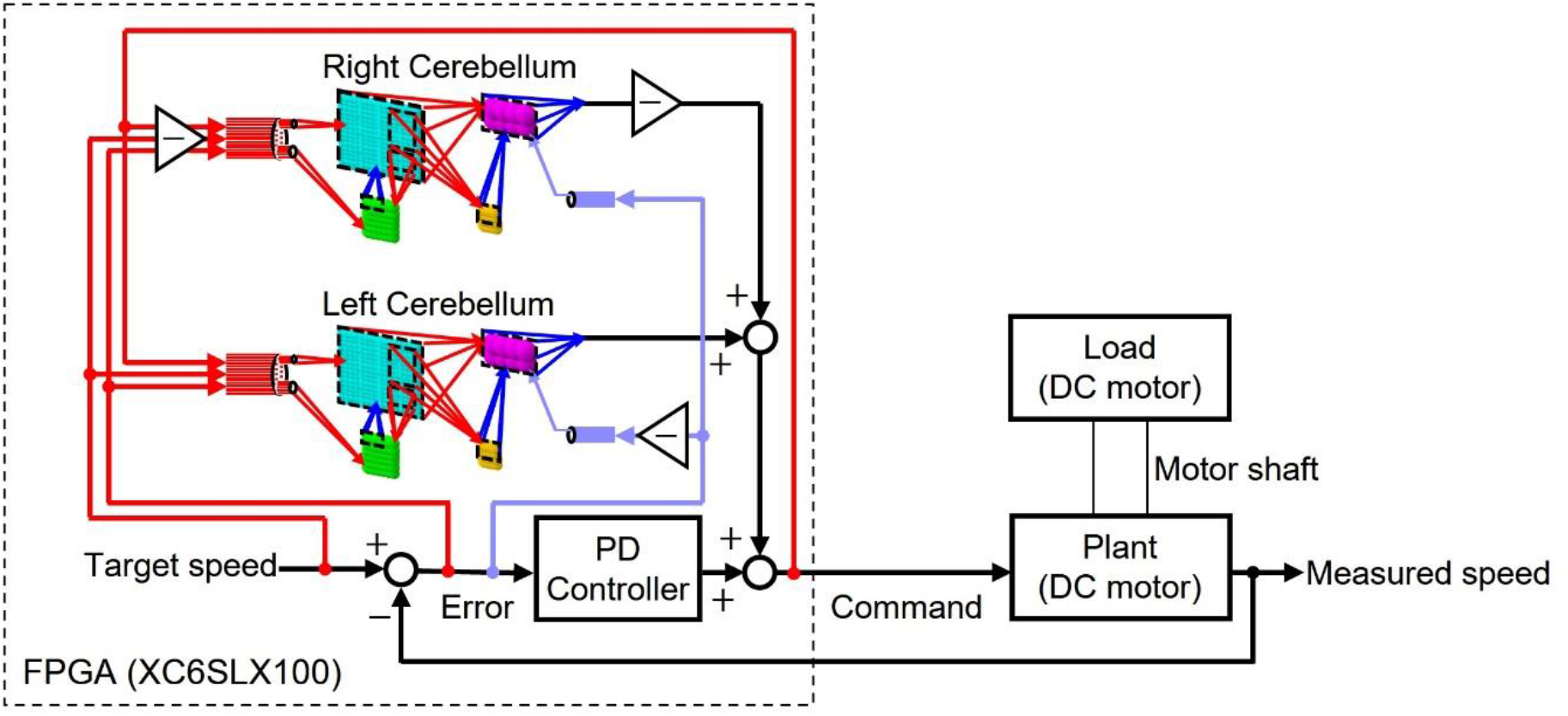
The structure of the control circuit in which the artificial cerebellum is implemented by FPGA. FPGA simulates two artificial cerebellums as left and right hemispheres. The target speed, the error and the command are input for MF of the artificial cerebellum, and error is input for CF. FPGA has a PD controller in parallel with the artificial cerebellum, imitating the neural circuit structure outside the cerebellum. The FPGA outputs the voltage as a command to the DC motor of the plant via the inverter by pulse width modulation. The rotation number of the DC motor is fed back to the FPGA by the encoder. The DC motor of Plant is connected to the DC motor of load by a shaft. When the load system is short-circuited, the Plant load fluctuates.

#### 2.2.2 Design of computation, communication, and memory systems

The differential equations describing the neuron model were discretized by the Euler method and implemented as digital circuits shown in Figure 3. Digital circuits that simulate neuron groups in the model are interconnected to form the cerebellar neural network shown in Figure 1. Figure 3A is the membrane potential processor (PP), B is the synaptic current processor (CP), C is the weight processor (WP), D is the synaptic conductance processor (DP), E is the synaptic plasticity processor (LP).

**Figure 3.**
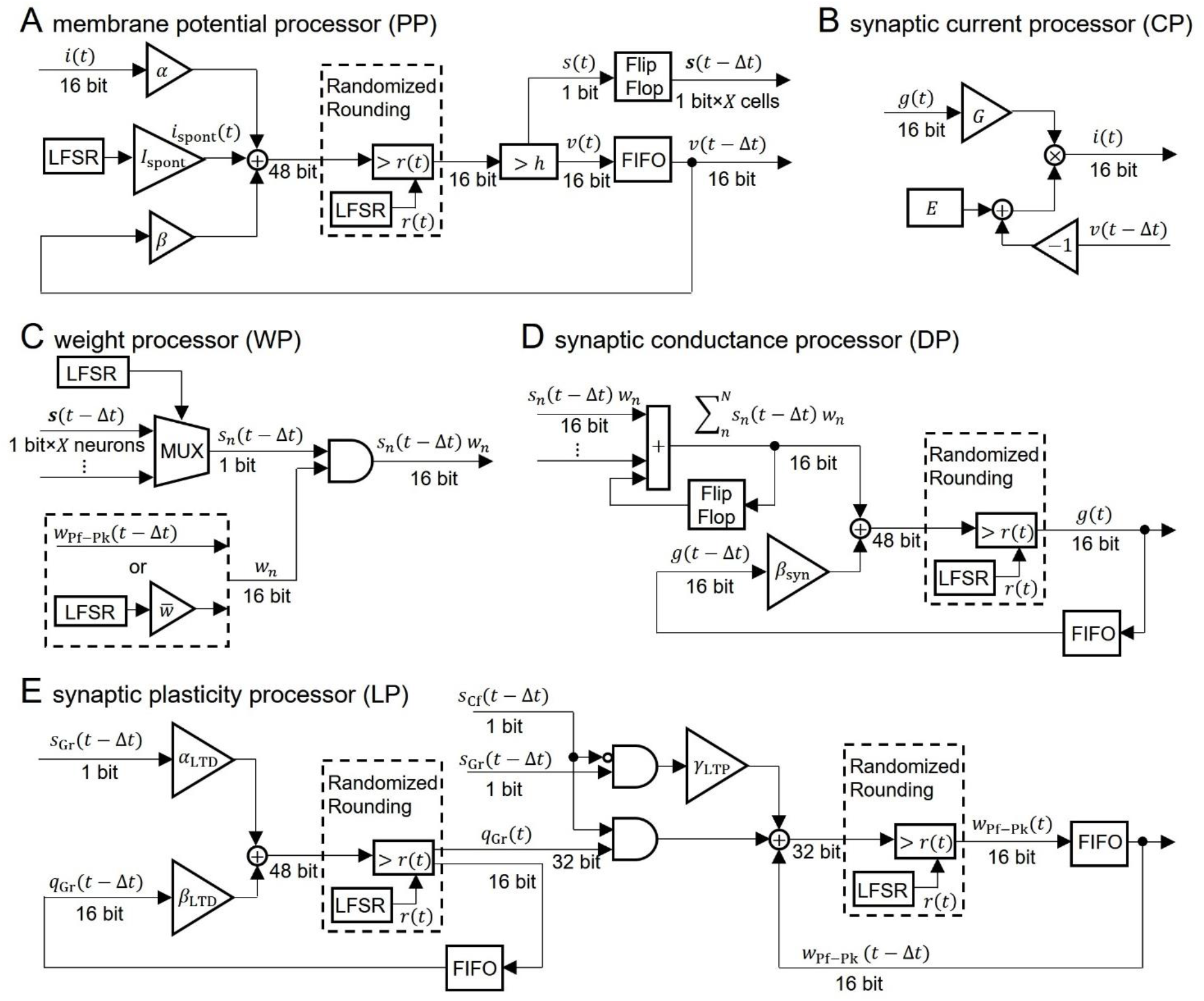
The block diagram of the neural processor. **(A)**, **(B)**, **(C)**, **(D)**, and **(E)** are processing devices for membrane potential, synaptic current, synaptic weighted, synaptic conductance, synaptic plasticity, respectively. The coefficients are defined as follows: *α* = 1/*C*, *β* = 1 – *g_L_*/*C*, *β_sym_* = 1 – 1/*τ_sym_*, *α_LTD_* = *γ_LTD_*/*τ_LTD_*, *β_LTD_* = 1 – 1/*τ_LTD_*. For the synapse transmission efficiency *w* in **(C)**, only the PF-PkC synapse uses the numerical value obtained by calculating LTD and LTP by **(E)**. Addition and AND are composed of logic circuits in the LUT. The DSP slice in the FPGA was used for the triangular block representing the gain and the multiplication. LFSR is a linear feedback shift register, each of which outputs a random number *r*(*t*) that differs depending on the initial seed. Flip Flop is a storage device that uses registers included in the look-up table (LUT) in the FPGA. The FIFO is a storage device that uses the block RAM in the FPGA in the first-in first-out format. The MUX is a multiplexer, and here, based on the LFSR signal, the spike events for one neuron are extracted from the flip-flop in which the spike events of some neurons (*X* neurons) are stored. Randomized rounding compares fraction bits with a random number in the LFSR, rounds up if the random number is smaller than the fraction, and rounds down if it is larger.

These processors adopted two parallel processing methods to complete the processing within a time step (1 ms). The first parallel processing method is pipeline processing used in all the processors in Figure 3. For example, pipeline processing in WP reduces the execution time by processing the second synapse at the MUX simultaneously with the stage of processing the first synapse at the AND. The second parallel processing method is the parallel operation of the dedicated processors described above. As shown in Figure 4, a number of PP, CP, DP, WP, and LP are provided for parallel processing. Since GrC has a large number of neurons, 4 each of processor are provided to use parallel processing. To process PkCs, 4 WPs and 4 LPs are provided because PkC has a large number of synapses despite a small number of neurons (8 neurons). 3 each of processors are provided to complete GoC processing during the PkC processing. one each of processors is provided, and BkC processing ends during PkC processing.

**Figure 4.**
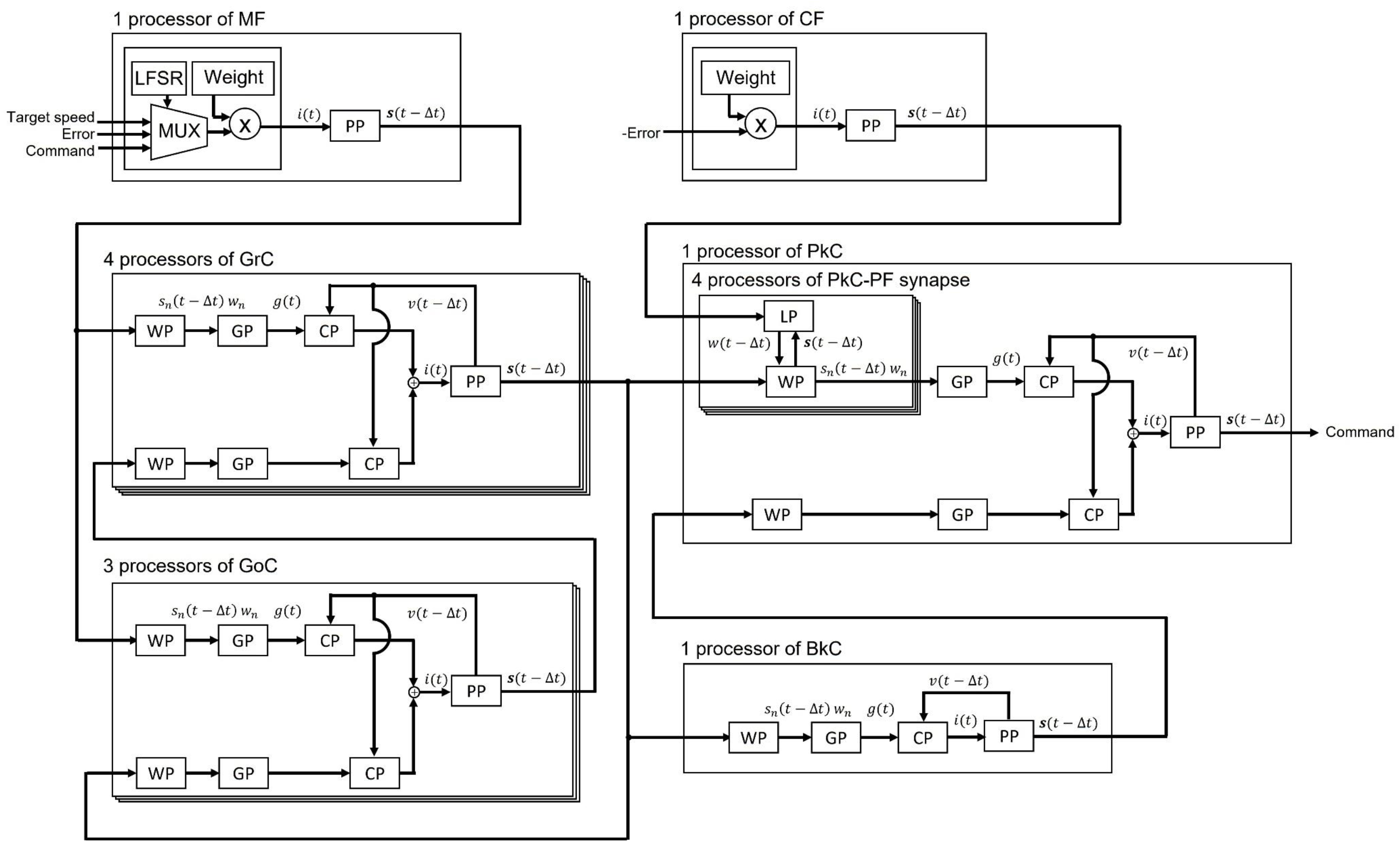
The processing circuit of the artificial cerebellum. Overlapping frames indicate that some processing circuits are provided.

In addition to these parallel processing, we adopted the following three new methods for implementing the artificial cerebellum efficiently on the FPGA within the limits of the number of logic blocks and time steps required for actual motor control.

##### 2.2.2.1 Fixed-point arithmetic with randomized rounding

In order to reduce computational cost and memory usage, 16-bit fixed-point numbers were adopted instead of floating-point numbers in the FPGA. However, round-off errors that occur in fixed-point arithmetic can degrade the precision of computation. In order to minimize the accumulation of rounding errors, random number rounding was adopted. This method compares a fraction that is supposed to be rounded up or off with random numbers. Namely, if the fraction is less than the random number, it is rounded up while if the fraction is greater than or equal to the random number, it is truncated. The average rounding of random numbers is distributed around the fractions and can be rounded unbiasedly. We employed uniform random numbers generated by a linear feedback shift register (LFSR). Although an LFSR is a pseudo-random number generator with periodicity, the bit width of the LFSR used in this study (32-bit) provides a sufficiently long period and can keep the bias in random number rounding small.

##### 2.2.2.2 Fully coupled spike transfer circuit

The memory of the spike event is frequently referenced by many processors. Short memory latency is required to take advantage of parallel processing. Therefore, we designed a parallel I/O interface composed of sender circuits and receiver circuits. We used this interface for the transmission of spike events between all neurons. An example of the interface is the transmission circuit from GrC to GoC as shown in Figure 5. The interface consists of sender circuits in the processor of GrC, which is the presynaptic neuron, and receiver circuits in the processor of GoC, which is the postsynaptic neuron.

**Figure 5.**
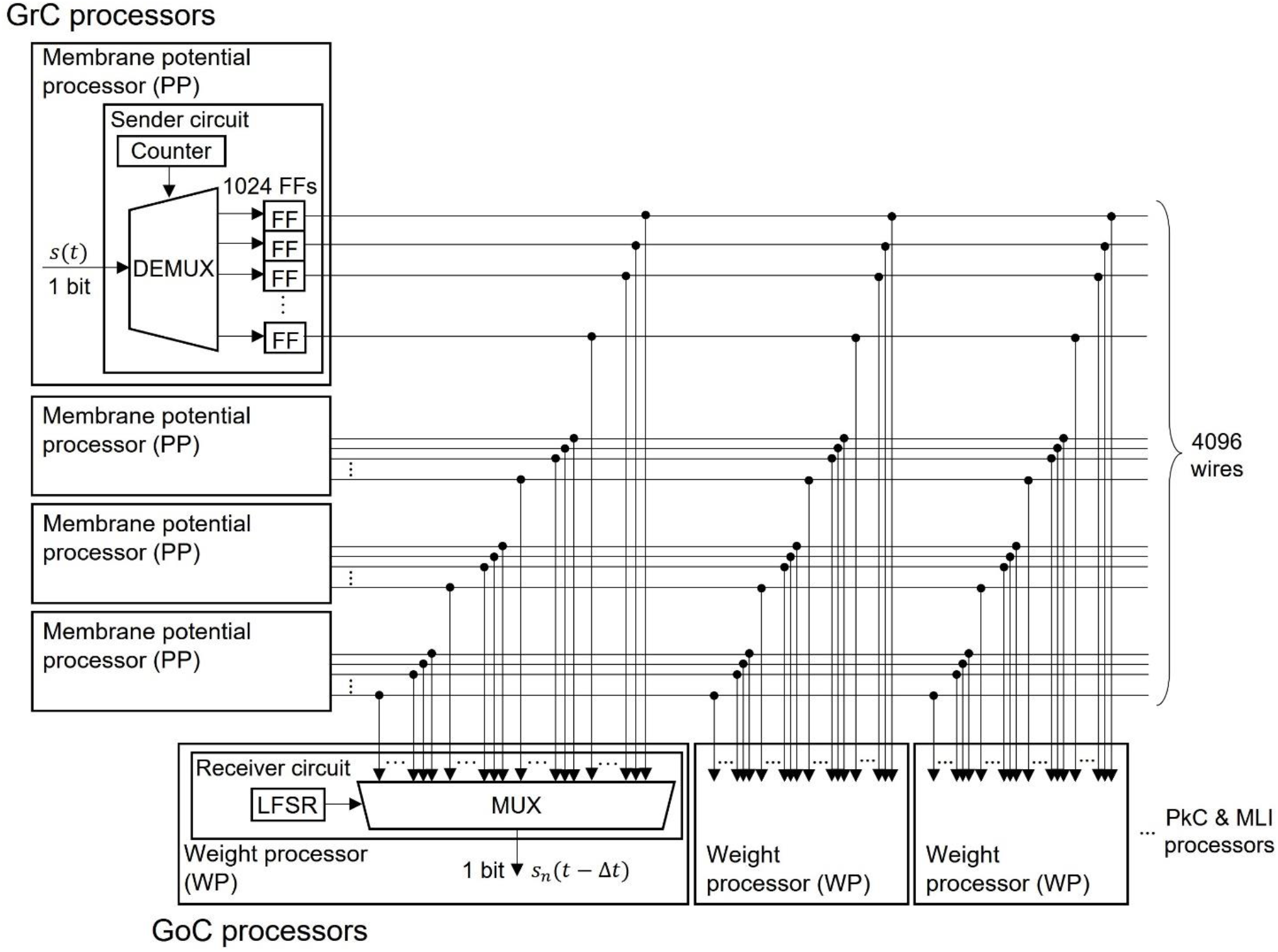
The structure of the fully coupled spike transfer circuit. The spike event *s*(*t*) calculated by the PP is stored in the flip-flop (FF). The outputs of all FFs of the presynaptic neurons are connected to MUX of all WPs of the postsynaptic neuron. By receiving the output of the LFSR, the MUX randomly selects the presynaptic neuron and outputs the spike event.

A sender circuit is composed of a counter, a demultiplexer (DEMUX) and flip-flops (FFs). The number of the FFs corresponds to the number of model neurons simulated in the PP of GrC; the number is 1024 in this case. When the membrane potential processing ends and the spike event is generated in the PP, the DEMUX receives the spike event and stores the event in one of the FFs depending on the counter output, which represents the neuron identifier (ID) of the spike event. The total number of spike events handled by this interface is 4096 because four sender circuits work in parallel. Here, the ON/OFF state of one wire of the FF output represents a spike event (whether spiked or not) of one neuron in a certain 1 ms.

A receiver circuit Is composed of an LFSR and a multiplexer (MUX), which selects one signal from its 4096 input signals depending on the LFSR output. The LFSR outputs pseudo random numbers, each of which represents the ID of the presynaptic neuron connected to the postsynaptic neuron processed at the period. The output of the LFSR is updated every 1 clock. One of the GoC (postsynaptic neuron) processors simulate 123 GoCs and three GoC processors simulate 369 GoCs in total. Each GoC is connected to 100 presynaptic GrC neurons, and the IDs of the connected GrC neurons are determined by the LFSR. Each receiver circuit repeats spike read procedure 100 × 123 times. When the circuit completes one cycle of the procedure, the LFSR is reset to the initial value which varies for each receiver circuit, and the spike event from the same presynaptic neuron ID is read in the next time step. By branching receiver circuits, the reading procedure of spike events is processed in parallel.

##### 2.2.2.3 Pseudo-random number generator to represent neural connections

In order to reduce memory usage, we adopted a pseudo-random number generator in place of a memory that stores information of neural connections. When implementing the artificial cerebellum on an FPGA, it is necessary to store neuron ID of presynaptic and postsynaptic neurons. As shown in Figure 3C, the ID is used by MUX to output spike events of a desired neuron group. The memory capacity required for storing the ID is the product of the number of pre-synaptic neurons, the number of post-synaptic neurons, and the convergence rate, which is huge (approx. 40 million bytes). However, the capacity of the internal RAM, which is the RAM embedded in an FPGA and provides much wider band width than the external RAM, is very limited. The effective use of the internal RAM is the key factor for implementing many neurons in an FPGA. A large amount of internal RAM should be used for storing differential equation variables, not for ID of neural connections. Because the neural connection in our model is defined by random numbers that are unchanged through an operation, we used an LFSR to achieve uniform random numbers that define neural connections.

Since an LFSR is composed only of XORs and FFs, internal RAMs are unnecessary. Similarly, synaptic weights that don’t have synaptic plasticity were generated by the LFSR.

## 3 Results

### 3.1 Specifications

The neural network constituting the artificial cerebellum contains 9,504 neurons (including MFs and CFs) with 240,484 synapses. The hardware resources used for the artificial cerebellum construction is shown in Table 3. RAMB16BWER and RAMB8BWER are 16-bit and 8-bit internal RAM, respectively. DSP48A1 is a DSP slice with a built-in adder and multiplier. At the FPGA clock frequency of 40 MHz, the calculation time for each hemisphere of the artificial cerebellum is 0.40 ms. The entire control circuit, including both hemispheres, completes all the computation in 1 ms time interval. Considering that the maximum firing rate of neurons in the cerebellum is about 500 spikes/s (Ito, 2012), 1 ms is fast enough for simulating the cerebellum. Therefore, real-time operation of the artificial cerebellum with this configuration is possible. The maximum power consumption at a clock frequency of 40 MHz and a device temperature of 25 °C was estimated to be 0.6 W by the Xilinx Power Estimator.

**Table 3.** Specifications of the artificial cerebellum

| Timing issues |  |  |  |
| --- | --- | --- | --- |
| Clock frequency |  |  | 40 MHz |
| Time step |  |  | 1 ms |
| Estimated Power Consumption in 25 °C |  |  |  |
| Maximum power consumption |  |  | 0.6 W |
| Typical power consumption |  |  | 0.4 W |
| Hardware resources utilization |  |  |  |
| Logic blocks | Number of useable | Number of used | Rate of utilization |
| Slice Registers | 126,576 | 24,402 | 19 % |
| Look Up Tables | 63,288 | 50,094 | 79 % |
| RAMB16BWER | 268 | 202 | 75 % |
| RAMB8BWER | 536 | 53 | 9 % |
| DSP48A1 | 180 | 160 | 88 % |

### 3.2 Simulation to evaluate the random number rounding

To evaluate the effect of rounding error, we calculated one neuron model with the following three methods: 1) Python with 64-bit floating point number, 2) Xilinx ISE Simulator with 16-bit fixed-point number without random number rounding, 3) Xilinx ISE Simulator with 16-bit fixed-point number with random number rounding. As a simulation with high calculation accuracy, we used Python’s float type 64-bits floating point number and performed the simulation on a PC.

We simulated one GrC receiving inputs from one GoC and one MF. The input signals sent from GoC and the MF are spike events whose firing time is determined by uniform random numbers generated by an LFSR. The simulation was performed under the condition that the GoC fires at an average of 31 spikes/s and the MF fires at an average of 62 spikes/s, so that the effects of rounding error can be easily evaluated. At this simulation, the weight between GrC and GoC was set to 93.8 pS, and that between GrC and MF was set to 320.0 pS.

Figure 6 shows the simulation results. The black lines in the figure plot the results simulated with 64- bits floating point number. Simulations with 16-bit fixed-point numbers cause rounding errors when computing the differential equations of synaptic conductance and membrane potential due to the limited number of digits. As the result, the synaptic conductance computed with 16-bit fixed-point numbers without random number rounding does not converge to 0 nS (Figure 6A, blue line), resulting in the positively biased membrane potential, and the increased spike frequency (Figure 6C, E, blue line and cross mark, respectively). In contrast, with randomized rounding, the synaptic conductance converges to 0 nS (Figure 6A, light cyan line), the membrane potential is not biased, and the frequency of spike occurrence is almost the same as that computed with 64-bit floating point numbers (Figure 6C, E, light cyan line and cross mark). The synaptic conductance computed with 64- bit floating point and that computed with 16-bit fixed point with random rounding are approximately equal. Even if an error occurs in the computation of synaptic conductance, the conductance computed with 16-bit fixed point with random number rounding converges to 0 nS while no spike comes to the neuron, so the accumulated error can be canceled. In the simulation computed with 64-bit floating point numbers, the mean firing rate of GrCs during 50 s was 6.58 spikes/s. The mean firing rate difference between the simulation with 64-bit floating point numbers and that with 16-bit fixed point numbers was 1.64 spikes/s. The mean firing rate difference between the simulation with 64-bit floating-point numbers and that with 16-bit fixed-point numbers with random rounding was 0.030 spikes/s. These results assure that the calculation accuracy can be maintained by using random number rounding when computing differential equations that describe a neuron model with fixed-point numbers.

**Figure 6.**
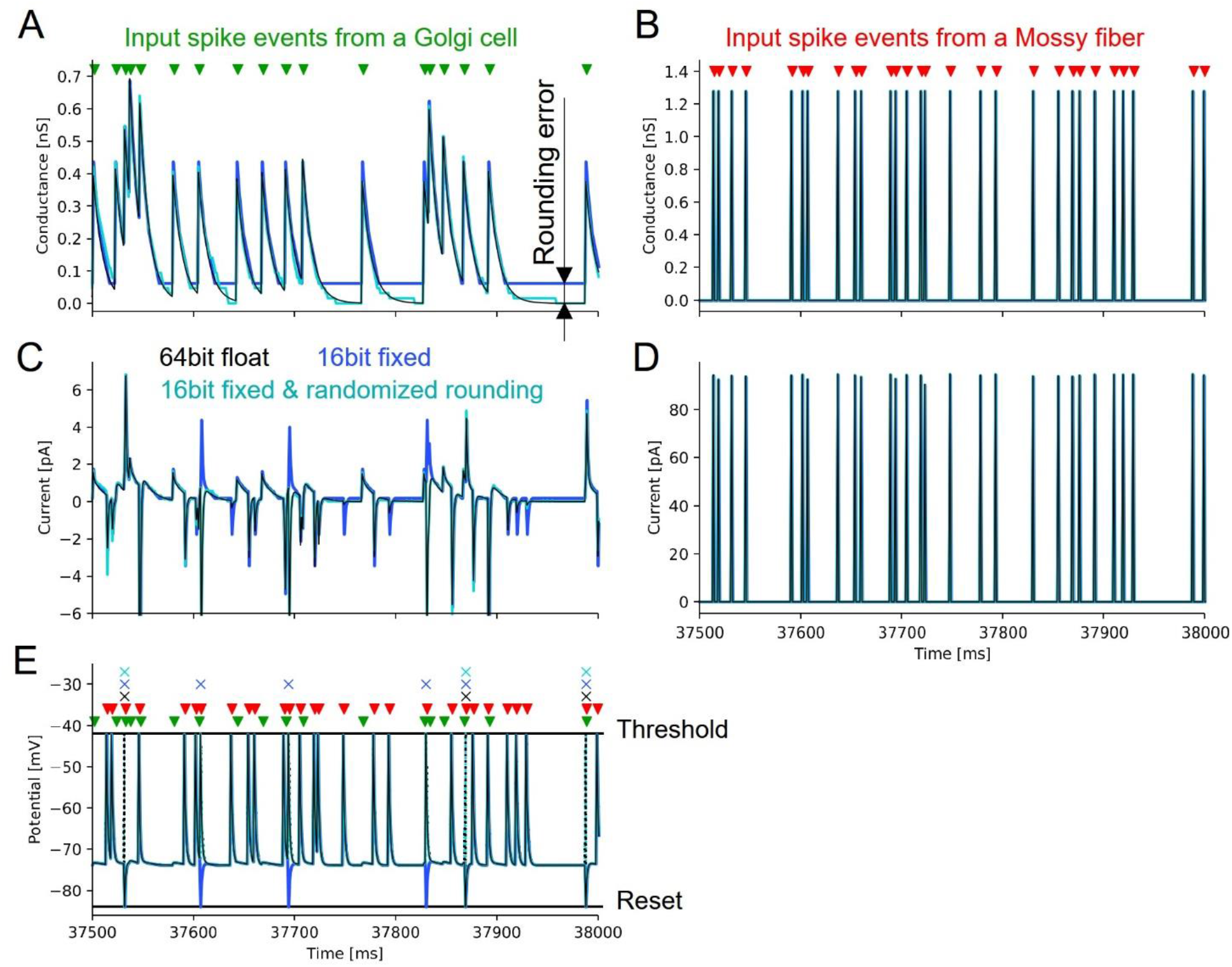
The simulation of the neuron model on VHDL and Python. The black line is the result of the simulation with rounding half up to 64-bit float by Python. Blue is the result of the simulation with rounding half up to 16-bit fixed point number by Xilinx ISE Simulator. Cyan is the result of the simulation with random number rounding to 16-bit fixed-point numbers by Xilinx ISE Simulator. Green triangles represent the spike timing of the input a GoC. Red triangles represent the spike timing of the input a MF. Cross marks represent the spike timing of the output a GrC. **(A)** is the postsynaptic conductance between the GrC and the GoC. **(B)** is the postsynaptic conductance between the GrC and the MF. **(C)** is the postsynaptic current between the GrC and the GoC. **(D)** is the postsynaptic current between the GrC and the MF. **(E)** is the membrane potential of the GrC. The dotted line in each simulation shows the change in membrane potential during spikes that is not stored in the FPGA.

### 3.3 Real-world Adaptive Machine Control

In order to evaluate the capability of the artificial cerebellum in adaptive actuator control in a real-world environment, we employed a DC motor and imposed a load whose strength changes in time. As shown in Figure 2, the FPGA controls the DC motor via an inverter in the control circuit. The rotation speed of the controlled object was fed back to the FPGA by the Hall effect sensor. The rotation speed error, which is calculated by subtracting the measured speed from the target speed, was input to the MFs and CFs of the artificial cerebellum.

Another DC motor was connected to the shaft of the controlled motor to impose a load. During the experiment, the switch in the circuit was opened to make the Load-OFF state and was closed to make the Load-On state. The on resistance of the switch was 13.5 Ω. The switch was opened and closed by the signal sent from the FPGA. The target time course of the rotation speed was a sine wave with an amplitude of 32 rotation per second (rps) and a period of 2.048 s. The load was turned on in the 120th cycle, and turned off in the 240th cycle, and turned on again in the 300th cycle, at which the control of the artificial cerebellum had been stable.

Figure 7 shows the results of motor control experiments repeated 10 times each with or without the artificial cerebellum. With the artificial cerebellum, the amount of error started from the same level as the PD controller alone. The adaptation did not start immediately due to the influence of noise in the real-world environment. The error started to decrease from around the 50th cycle due to motor learning in the artificial cerebellum. When a load was imposed in the 120th cycle, the error rose sharply, but gradually decreased as the artificial cerebellum adapted to the load. By contrast the error increased with much smaller amount at the timing of Load-Off in the 240th cycle and Load-On in the 300th cycle, demonstrating generalization in adaptation to both Load-Off and Load-On states.

**Figure 7.**
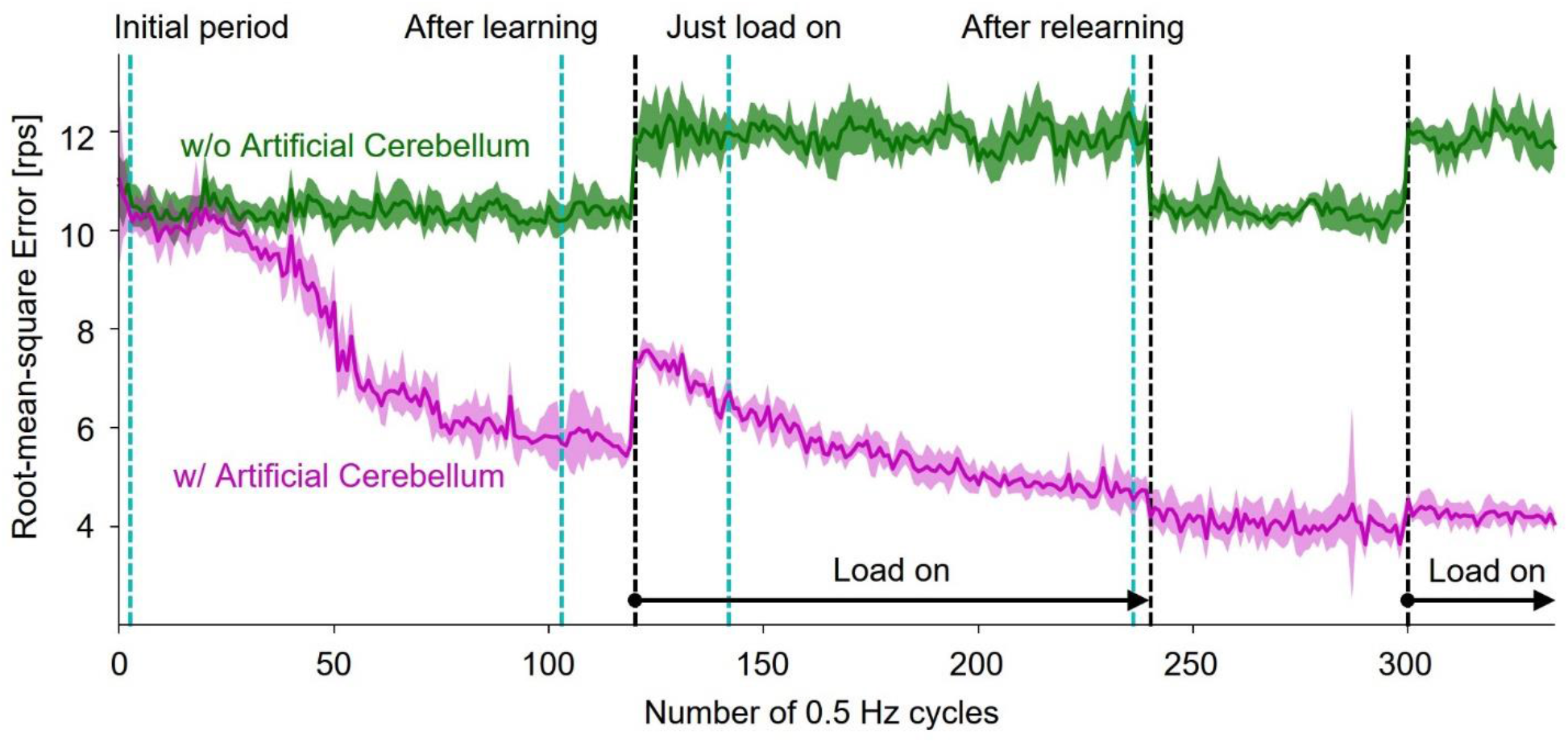
The adaptive motor control by the artificial cerebellum in the real world. The magenta line is the control result with the artificial cerebellum. The green line is the result of controlling with the PD controller without the artificial cerebellum. The solid line and fill represent the mean and the standard deviation of 10 experiments. The black dashed line represents the Load on section. The cyan dashed line represents the number of periods evaluated in Figure 8.

Figure 8 shows rotation speed of the controlled DC motor and the activities of representative neuron models in the following four periods: initial period, after learning, just load-on, and after relearning corresponding to each triangle marked on the horizontal axis in Figure 7. In the top panels of Figure 8, the black, green, and magenta lines plot the target speed, the measured speed controlled without the artificial cerebellum, and the measured speed controlled with the artificial cerebellum, respectively. In the initial period (left top panel), the measured speed controlled with artificial cerebellum (magenta) was smaller than and delayed to the target speed (black). This situation was similar to the speed controlled without artificial cerebellum (green). After learning (2^nd^ column from the left, top panel), the measured speed controlled with artificial cerebellum (magenta) improved the amplitude and the delay. After load on (2^nd^ to the right column, top panel), the speed controlled with artificial cerebellum (magenta) decelerated just before it reached to the maximum speed. After relearning (right top panel), the control with artificial cerebellum (magenta) improved the deceleration just before it reached to the maximum speed, and further improved the delay after switching the direction of rotation. The raster plots in the 2nd to the bottom panels of Figure 8 show spike timings of 8 neurons randomly selected from each neuron type in the left (red) and right (blue) cerebellar hemispheres. The red and blue lines in these panels are average firing rate of the 8 neurons in left and right cerebellar hemisphere, respectively. MF and CF firing rates (2^nd^ and 3^rd^ row, respectively) showed almost the same responses in all cycles. The firing rate of GrC (3^rd^ row from the top) decreased significantly in the after-learning period comparing to those in the initial period. The same trend is observed in the relationship between GrC firing rates in the after-relearning period and those in the just-load-on period. The firing rates of GoC (3^rd^ row from the bottom) and MLI (1^st^ row from the bottom) showed similar changes to those of GrC. Because the initial weights of the synapse between PF and PkC was 0, the firing rate of PkC (2^nd^ row from the bottom) showed only spontaneous firing rate in the initial period. In the after-learning period, the firing rate of PkC increased mainly after switching the direction of rotation. In the just-load-on period, the firing rate of PkC was active and was similar to that in the after-learning period. In the after-relearning period, the firing rate of PkC was further increased after switching the direction of rotation. These results assure that the FPGA artificial cerebellum may provide useful information as to signal processing executed in the cerebellar cortical neural network consisting of these neuron types connected in a manner unique to the cerebellum while working as an adaptive motor controller in real-world.

**Figure 8.**
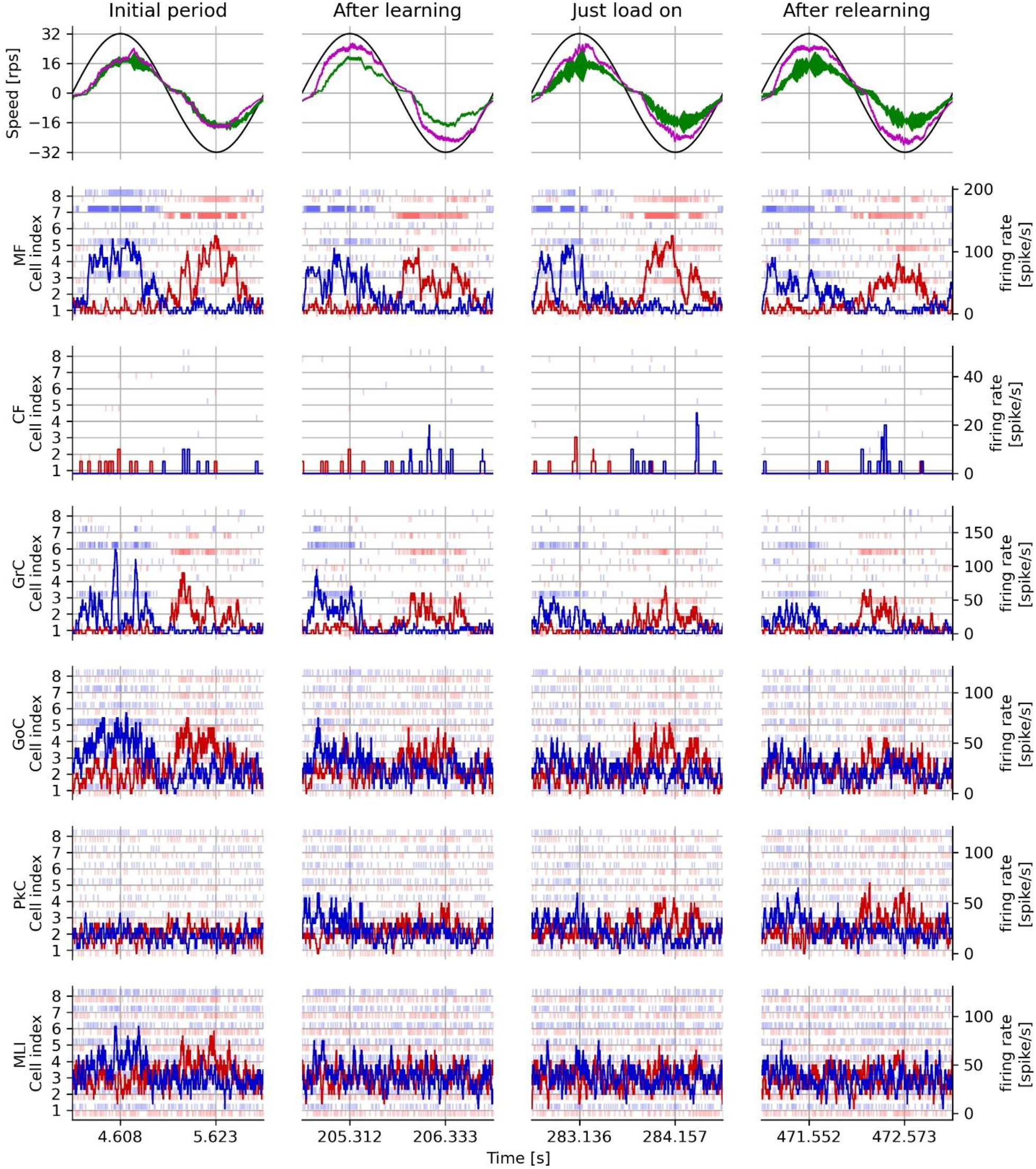
The measured rotation speed and activities for each neuron type in adaptive motor control. From the left column, the initial period, the period after learning, the period after load on, and the period after relearning with the load on in one experiment are shown. The first row from the top shows the target speed (black line), the measured speed without artificial cerebellum (green line), and the measured speed with artificial cerebellum (magenta line). The second and subsequent rows from the top show the responses of each type of neuron in the left cerebellum (blue) and right cerebellum (red). The light vertical line markers indicate spike timing of 8 neurons, and dark solid lines indicate average firing rate of 8 neurons.

## 4 Discussion

We implemented for the first time a spiking artificial cerebellum on an FPGA, which accurately replicated the structure of cerebellar neural circuitry, including MLIs, and runs in real time. We evaluated calculating the neuron model (Figure 4) and demonstrated its capability as a real-time adaptive controller in the real-world experiments (Figure 5 and 6). In the following, we discuss general advantages of FPGAs as a hardware platform for the implementation of neural network models of the spiking artificial cerebellum. Further, we discuss three key techniques used in the current FPGA implementation of the artificial cerebellum.

### 4.1 FPGA as a hardware for implementing the spiking artificial cerebellum

There are currently two major hardware platforms that support efficient implementation of neural networks. One is analog circuits represented by Neurogrid (Benjamin et al., 2014), and the other is non-Neumann type custom chips such as TrueNorth (Merolla et al., 2014), SpiNNaker (Furber et al., 2013), and programmable logic devices including FPGAs. The former has problems of instability in execution and difficulty to reconstruct the hardware design. Thus, it would be difficult to implement LTD/LTP of Purkinje cell, other synaptic plasticity discovered (Gao et al., 2012; Mapelli et al., 2022; Bonnan et al., 2023) and gap junctions (Ito, 2012; De Zeeuw et al., 2021) in the cerebellum. In addition, they are generally specialized for random networks or deep neural networks and are not necessarily the efficient implementation method for shallow neural networks such as that of the cerebellum. By contrast, the latter digital chips including FPGAs are programable and have generally higher noise immunity than analog circuits.

So far, real-time simulations of spiking artificial cerebellum have been performed using GPUs (Yamazaki and Igarashi, 2013; Florimbi et al., 2021; Kuriyama et al., 2021), supercomputers (Yamazaki et al., 2019; Yamaura et al., 2020), and FPGAs (Luo et al., 2016; Xu et al., 2018) It is shown that FPGAs have a clear advantage especially in power consumption. The maximum power consumption of our FPGA artificial cerebellum was 0.6 W in total, and 61 uW/neuron without considering the number of synapses and neuron types. This number is far much less than those in other artificial cerebellums simulated in the K computer (185 uW/neuron, (Yamaura et al., 2020) and in GPUs (13,383 uW/neuron, (Kuriyama et al., 2021).

Furthermore, FPGA design with VHDL can also be used for ASIC implementation, which can eliminate unused elements and redundant wiring. It has been shown that ASIC can reduce semiconductor area to 1/21, delay to 1/2.1, and power consumption to 1/9.0 compared to FPGA in 90 nm process (Kuon and Rose, 2006). Although the FPGA used in the current study was a spartan-6 with a 45 nm process and cannot be directly compared, it may be possible to achieve 65 mW when implemented on an ASIC. This is below 100 mW (Kim et al., 2007), at which semiconductor devices would thermally damage living organisms.

Another critical difference between FPGA platforms and processor-based implementation is the band width between processors and memories. FPGAs can provide much wider band width because they have many distributed localized memory banks each of which can be accessed exclusively from dedicated processors synthesized in the FPGA. Therefore, FPGA platform can avoid memory wait, which often occurs in systems with shared global memory resources accessed from many processors.

### 4.2 Three techniques to implement the artificial cerebellum on the FPGA

#### 5.2.1 Fixed-point arithmetic with random rounding

Computing numerical solutions of differential equations in fixed-point numbers produces a constant rounding error unless random rounding is used (Figure 4). This type of rounding error is commonly encountered in first-order lag differential equations describing the synapse model and the neuron model. The first-order lag differential equation is expressed by the following equation:

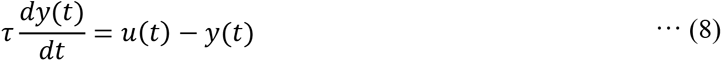

where *u*(*t*) is the input, *y*(*t*) is the output, and *τ* is the time constant. The formula for rounding half up the binary number of arbitrary digits *n* of the first-order lag differential equation discretized by the Euler method can be expressed by the following equation:

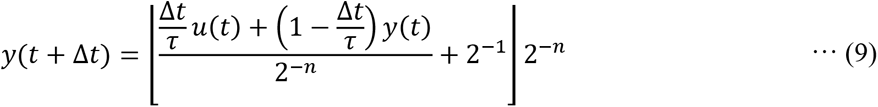

where |*x*| is the floor function. *n* is the number of digits representing fractional part of *y*(*t*). When the input *u*(*t*) is 0 and the output *y*(*t*) is smaller than the following particular value, *y*(*t*) ceases to change due to a constant rounding error. This condition is −2^−*n*−1^ *τ*/Δ*t* ≤ *y*(*t*) < 2^−*n*−1^ *τ*/Δ*t*. This means that the longer the time constant (the closer the coefficient *τ* is to 1), the larger the range of rounding errors. For example, let the coefficient *τ*/Δ*t* is 2^5^ and fixed-point numbers of (1 – Δ*t*/*τ*) and *y*(*t*) have the 16-bit fractional part. The fractional part of the multiplying (1 – Δ*t*/*τ*) and *y*(*t*) is 32 bits, so it must be rounded to 16 bits (= *n*). At this time, the term that the constant rounding error remains is −2^−12^ ≤ *y*(*t*) < 2^−12^, which is within 16 bits of significant digits of *y*(*t*). In this way, if the output *y*(*t*) matches the condition described above, it is necessary to perform random number rounding. In particular, when computing second-order integration in the synapse model and the neuron model, random rounding should be adopted because the bias is occurred by the constant rounding error in the first step and the constant rounding error accumulates in the second step.

Thus far, 16-bit fixed-point numbers and random rounding have been adopted to streamline deep learning operations, and have shown accuracy comparable to 32-bit floating-point numbers (Gupta et al., 2015). In FPGA spiking artificial cerebellum, we showed that 16-bit fixed-point numbers and random number rounding can compute spikes with the comparable accuracy to 64-bit floating-point numbers. Hence, fixed-point arithmetic with random rounding is capable of efficient implementations while preserving arithmetic accuracy.

#### 5.2.2 Fully coupled spike transfer circuit

In order to achieve efficient communication of spike events, we designed a fully coupled spike transfer circuit, which is a parallel input/output interface. The strengths of using this interface for FPGA implementation of the artificial cerebellum include the following points.

First, it reduces the time required for spike transfer. Since the GrC in the cerebellum has parallel fibers connecting to a large number and variety of neurons over a wide area, spike event memories are accessed at very high frequency. In our artificial cerebellum, 8 WPs simultaneously access the spike event memories at 40 MHz. If the event is stored in a memory that is accessible only in serial, the time required to write/read the spike event increases as a multiplier of the number of presynaptic and postsynaptic neuron processors. In this interface, All the spike events are accessible in parallel from the processors that receive the spike events because all the spike events are stored in the FFs whose outputs are accessible from any circuits in the FPGA. In addition, each read and write procedure completes in 1 clock, even if the number of neurons is large and there are many processing circuits for writing and reading. This interface reduces the spike transfer time drastically.

Second, the spiking neuron model and the FPGA are suitable for implementing the fully coupled spike transfer circuit. In this interface, the numbers of the wires and the input ports of the MUX increase as the number of neurons increases. However, the increase in the circuit scale can be kept small because information transferred in this interface is the spike event, which can be expressed with only 1 bit. On the other hand, FPGA essentially has many lookup tables including logic circuits and many wires (the FPGA used in this study has 63,288 lookup tables with 380,000 wires).

Therefore, the interface for a larger number of presynaptic/postsynaptic neurons is installable in an FPGA.

#### 5.2.3 Pseudo-random number generator to represent neural connections

As previously stated in Section 3.2, the utilization of an LFSR as a pseudo-random number generator serves to conserve memory. However, it should be noted that these pseudo-random numbers can have an impact on the structure of the cerebellar network.

The convergence was established as a fixed number of mean values obtained through autopsies. However, due to the uniform distribution and overlap of the pseudo-random numbers determining which presynaptic neuron to connect to, it is possible for the same presynaptic neuron to be selected several times. As the processing of synapse formation is equal when the same presynaptic neuron is chosen, this is equivalent to doubling the weight and decreasing the convergence by one. When the weight is doubled, the initial value is determined by a random number, leading to potential bias in rare cases.

Divergence is the sum of some uniformly distributed pseudo-random numbers. The range of these numbers is [*a*,*b*] = [1,number of presynaptic neurons – 1], with a mean of *μ =(a+b)/2* and a variance of *σ* = (*b* – *a*)^2^/12. When the number of uniform random numbers (= *n*) is sufficiently large, the central limit theorem states that divergence follows a normal distribution with parameters *N*(*nμ*,*nσ*/12) and lies within the range [*an,bn*]. However, it should be noted that the convergence and divergence present in the cerebellum, as observed through anatomy, are not constant (Eccles et al., 1967; Ito, 2012), hence the utilization of pseudo-random numbers in this model. Additionally, as each Purkinje cell receives connections from more than 100,000 GrCs, this model does not use random numbers and instead assumes projections from all GrCs.

Furthermore, each mossy fiber branches to form 20-30 presynaptic sites and inputs to various GrCs and GoCs (Eccles et al., 1967). The axons of GrCs project as parallel fibers over a distance of approximately 2 mm in the cerebellum (Eccles et al., 1967), while the maximum linear extent of GoC axons is 650 ±179 μm in the sagittal plane and 180 ±40 micrometers in the medial-lateral plane (Barmack and Yakhnitsa, 2008). Astrocyte axons in the molecular layer of the cerebellum are approximately 200 micrometers in diameter (Barmack and Yakhnitsa, 2008). Based on a reported density of granule cells of 4x10^6^ mm^3^ (Solinas, 2010), the granule cell layer of this model, containing 4096 GrCs per hemisphere, corresponds to a cubic volume of 100 micrometers on each side. As the axon of each neuron is long enough to span the volume of the model, each neuron in this model was selected as a presynaptic neuron at random.

## 5 Summary and Conclusions

We implemented an artificial cerebellum constructed with a spiking neural network on FPGA based on known anatomy and physiology. The artificial cerebellum is composed of the left and right hemispheres including 9,504 neurons and 240,484 synapses representing all the major cerebellar cortical neuron types and their synaptic connections. The FPGA artificial cerebellum is compact (< 54x86 mm), lightweight (< 32 g), low power consumption (< 0.6 W), and runs in real time (1 ms intervals). In real-world experiments, the FPGA artificial cerebellum was demonstrated to function as an adaptive motor controller in which spiking activities of selected neurons can be monitored to analyze neural mechanisms of cerebellar motor learning. With these specs and capabilities, we conclude that the FPGA artificial cerebellum currently configured has the potential to be utilized in various adaptive motor control problems while allowing us to evaluate information processing executed in the cerebellar neuronal circuitry in real-world environment.

## 6 Conflict of Interest

The authors declare that the research was conducted in the absence of any commercial or financial relationships that could be construed as a potential conflict of interest.

## 7 Author Contributions

YS conducted hardware implementation and all the simulations and real-world simulations. YS, HO, and YH wrote and edited the manuscript. All authors contributed to the article and approved the submitted version.

## 8 Funding

JSPS KAKENHI Grant Number 20H04286, 19K06756, 18KK0286, and JST CREST Grant Number JPMJCR22P5 to YH. JSPS KAKENHI Grant Number 19K12916 to HO.

## 9 Abbreviations

FPGA, field-programmable gate array; MF, mossy fiber; CF, climbing fiber; GrC, granule cell; GoC, Golgi cell; BkC, basket cell; PkC, Purkinje cell; PF, parallel fiber; LTD, long-term depression; LTP, long-term potentiation; FF, flip flop; MUX, multiplexer; DEMUX, demultiplexer; LFSR, linear feedback shift register; PD controller, proportional–derivative controller; PP, membrane potential processor; CP, synaptic current processor; WP, weighted processor; DP, synaptic conductance processor; LP, plastic processor; ID, identifier.

## Acknowledgments

The authors would like to thank Profs. Minoru Asada and Ichiro Tsuda for their valuable comments on the preliminary results of our real-world motor control experiments.

## Notes

### Competing Interest Statement

The authors have declared no competing interest.

